# Identification of a key residue in the cellular transcription factor BCL11b important for its global acetylation and its nuclear localization

**DOI:** 10.64898/2026.01.19.700445

**Authors:** Laure Vreux, Caroline Vanhulle, Mathilde Galais, Sylvain Fauquenoy, Estelle Plant, Thomas Loustau, Maxime Bellefroid, Gwenaëlle Robette, Maryam Bendoumou, Marion Santangelo, Valérie Martinelli, Christian Schwartz, Ruddy Wattiez, David Communi, Olivier Rohr, Carine Van Lint

## Abstract

The cellular transcription factor BCL11b (B-cell CLL/lymphoma 11b) interacts with numerous cellular and viral factors to modulate gene expression positively or negatively. Post-translational modifications of BCL11b, such as SUMOylation and phosphorylation, have been documented to switch its transcriptional activity from a repressor to an activator state. In the present study, we investigated the acetylation of BCL11b and we identified the histone acetyltransferase p300 as able to acetylate BCL11b. Subsequently, we observed that the mutation of the lysine K686 residue of BCL11b (BCL11b K686R) influenced its global acetylation. Furthermore, the BCL11b K686R mutation also modulated the transcriptional regulation of BCL11b, including its activity in regulating the p21 and IL-2 promoters. This effect on transcriptional regulation was due to the importance of the lysine K686 residue for BCL11b nuclear localization. Our results underscore the critical role of the lysine K686 residue in BCL11b for its interaction with p300 and its nuclear localization, suggesting a possible function of p300 in the nuclear transport of BCL11b. Collectively, our findings contribute to a better understanding of BCL11b-mediated gene expression and of the interactions of BCL11b with cellular partners.

## Introduction

BCL11b (B-cell CLL/lymphoma 11b), also known as CTIP2 (COUP-TF Interacting Factor 2), is a zinc finger transcriptional co-factor involved in neural development and lymphomagenesis^1,2^. Functionally, BCL11b can act as an activator or repressor of transcription depending on the associated-factors and the epigenetic modifications it causes^3–9^. As a transcriptional repressor, BCL11b recruits the NuRD (Nucleosome Remodeling and Deacetylase) complex, which functions by inducing heterochromatin formation through deacetylation of histones by histone desacetylases (HDAC). This results in the suppression of various cellular genes including *Cd4* and *Hes1*^10,11^. Furthermore, we and others have previously demonstrated that BCL11B and NuRD complexes are recruited to the HIV-1 promoter, resulting in repression of basal and Tat-transactivated viral transcription. We have also demonstrated that BCL11b is associated with the inactive P-TEFb (positive transcription elongation factor b) complex through interaction with HEXIM1 and loop 2 of the 7SK snRNA^4–6,12–14^. In microglial cells, BCL11b collaborates with HMGA1 (non-histone chromatin protein High mobility group AT-hook 1) to repress P-TEFb-dependent genes, including the HIV-1 promoter^13^. Depletion of BCL11b and HMGA1 leads to an increase in the initiation phase and the Tat-dependent phase of HIV-1 transcription. HMGA1 facilitates the recruitment to the HIV-1 promoter of BCL11b associated with the 7SK snRNA complex, leading to the repression of HIV-1 transcription^13^. In microglial cells, BCL11b is recruited to the Sp1 binding sites of the cellular CDKN1A/p21^WAF^ promoter and of the viral HIV-1 promoter, thereby inducing their repression. Once recruited to Sp1 binding sites, BCL11b recruits chromatin remodelers such as HDAC1/2 and the histone methyltransferase SUV39H1 to the promoter region, inducing epigenetic modifications and the recruitment of Heterochromatin Protein 1 (HP1)^3,6^.

Despite being widely characterized as a transcriptional repressor, BCL11b can also function as a transcriptional activator due to the dynamic regulation of its post-translational modifications (PTMs) such as phosphorylation and SUMOylation^1^. Phosphorylation involves the covalent addition of a phosphate group to specific amino acids, primarily serine or/and threonine residues, leading to changes in the structural conformation of the protein or in the creation of a protein anchor point^16^. SUMOylation PTM involves the covalent addition of SUMO peptide(s) on lysine residue(s) and this PTM is known to regulate the activity of transcription factors by affecting their cellular localization, stability and/or interactions with other proteins^17^.

The PTMs of BCL11b, particularly SUMOylation and phosphorylation, play essential roles in regulating T-cell development^15,18–20^. In unstimulated CD4^+^ T cells, BCL11b is associated with the NuRD complex and interacts directly with metastasis-associated protein MTA1, resulting in the repression of the IL-2/Id-2 promoter. However, upon T-cell receptor activation, the serine S2 of BCL11b is phosphorylated through the Mitogen-activated protein kinase (MAPK) pathway, leading to the disruption of its interaction with MTA1. Consequently, the lysine K689 of BCL11b undergoes SUMOylation, allowing the recruitment of p300 to activate transcription^15,18^. After 2-5h of PMA/ionomycine treatment, the cellular transcription factor KLF4 leads to downregulation of *BCL11b* gene expression, resulting in decreased activation of IL-2 and Id-2 promoters^15,18^.

Vogel and colleagues have identified 50 different phosphorylation sites on the human version of BCL11b and 37 phosphorylation sites on the murine version of BCL11b^19^. Phosphorylation and SUMOylation are crucial for regulating the transcriptional activity of a promoter targeted by BLC11b^19^. However, acetylation of BCL11b has not yet been explored, even though it is a significant PTM that promotes conformational changes and controls protein-protein interactions. Additionally acetylation has been shown to regulate the subcellular localization of several proteins^21,22^.

In the present study, we characterized the role of the histone acetyltransferase p300 in BCL11b acetylation using immunoprecipitation experiments. We revealed that lysine K686 residue in BCL11b is critical for its global acetylation. Importantly, we also demonstrated that K686 is indispensable for the transcriptional regulation of BCL11b as well as for its nuclear localization. p300 interacted with BCL11b through K686, and depletion of p300 led to cytoplasmic localization of wild-type BCL11b. These observations suggest that p300 may play a role in the shuttling of BCL11b between the cytoplasm and the nucleus. Overall, our study highlights the functional significance of a single amino acid residue in BCL11b, which impacts its interactions with other cellular partners and ultimately its biological function.

## Results

### BCL11b is acetylated by the acetyltransferase p300

So far, a few studies have focused on the post-translational modifications of BCL11b^15,18,19^. Acetylation of BCL11b has never been reported in the literature. p300 (histone acetyltransferase p300), pCAF (p300/CBP-associated factor) and CBP (CREB-binding protein) are transcriptional cofactors known to have acetyltransferase activity by adding acetyl group to transcription factor and histone lysines^23^.

To investigate potential acetylated forms of BCL11b, we overexpressed a FLAG-tagged version of BCL11b with different known acetyltransferases (i.e the wild-type p300, pCAF and CBP acetyltransferases or the HAT-deleted (ΔHAT) mutant of these three acetyltransferases) in the human embryonic kidney HEK293T cell line. We found that p300 was the only acetyltransferase inducing an acetylated form of BCL11b, as shown by immunoprecipitation and Western blot experiments using an anti-acetyl antibody. In contrast, overexpression of a mutated form of p300 lacking the acetyltransferase domain (ΔHAT) did not induce acetylation of BCL11b (**Fig. 1A**). Our findings demonstrate that BCL11b can be acetylated and that the acetyltransferase p300 could be responsible for this PTM.

**Figure 1:**
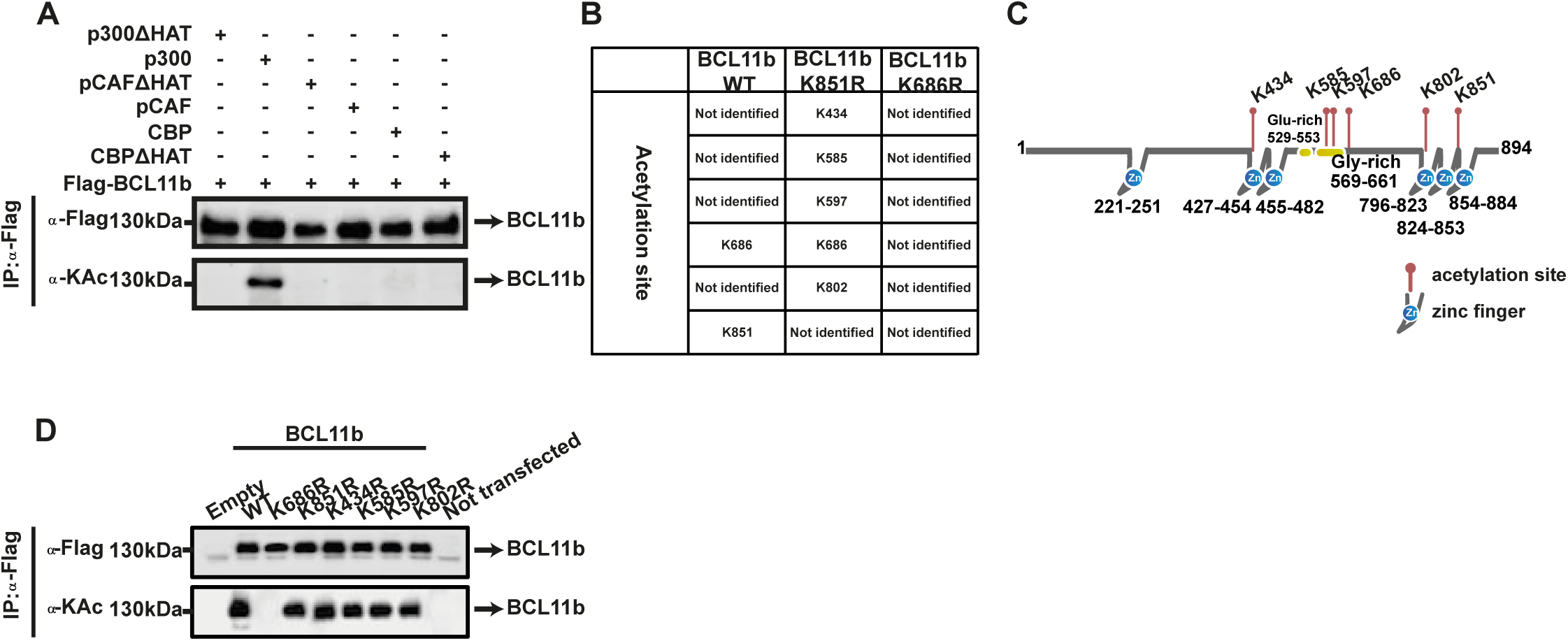
BCL11b is acetylated on six different lysine residues by the histone acetyltransferase p300. **(A)** HEK293T cells were transfected with an expression vector for wild-type BCL11b and with an expression vector for either the wild-type p300, either the wild-type p/CAF, or the wild-type CBP or with an expression vector for either a p300 depleted in its HAT domain, either a p/CAF depleted in its HAT domain, or a CBP depleted in its HAT domain. After forty-eight hours post-transfection, the total protein extracts were collected and immunoprecipitated with an antibody directed against Flag-BCL11b or with IgG as negative control. The eluted protein complexes were analyzed by Western blot using specific antibodies against BCL11b or against acetylated lysines (KAc). **(B)** HEK293T cells were transfected with expression vector for p300, and with expression vector for either BCL11b wild-type or mutated (K686R or K851R) BCL11b. After forty-eight hours post-transfection, the total protein extracts were collected and immunoprecipitated with an antibody against Flag-BCL11b or with IgG as negative control. The eluted protein complexes were analyzed by mass spectrometry to identify the acetylated lysines. Mass spectrometry results are shown in Table B. **(C)** Schematic representation of the BCL11b protein with the localization of the acetylated lysines we identified by mass spectrometry. **(D)** HEK293T cells were transfected with expression vectors for p300, BCL11b wild-type or mutated (K686R, K851R, K434R, K585R, K597R and K802R). After forty-eight hours post-transfection, the total protein extracts were collected and immunoprecipitated with an antibody against Flag-BCL11b or with IgG as negative control. The eluted protein complexes were analyzed by Western blot using specific antibodies against BCL11b or against acetylated lysines (KAc).

### Lysine K686 of BCL11b is essential for its global acetylation

We next performed mass spectrometry analysis to identify the lysine residues in BCL11b that were acetylated in the presence of p300 overexpression. We identified two acetylated lysine residues, K686 and K851 in BCL11b. In order to study the role of these two lysines residues on BCL11b global acetylation, we performed site-directed mutagenesis to substitute both lysines by an arginine which mimics a constitutively non-acetylated amino acid (BCL11b mutants K686R and K851R, respectively) (**Fig. 1B**). Mass spectrometry analysis of mutant BCL11b K851R identified in addition to K686 four new acetylated lysines (K434, K585, K597, K802) that were not acetylated in wild-type BCL11b. However, no acetylated lysine residues were identified in the mutant BCL11b K686R (**Fig. 1B**). All acetylated lysines of BCL11b are represented in **Figure 1C**. Site-directed mutagenesis was then performed to study the impact of the four additional lysines on BCL11b global acetylation. Immunoprecipitation targeting BCL11b followed by western blot targeting acetylated lysines demonstrated that mutation of lysine K686 impacted the global acetylation of BCL11b, whereas mutation in the other lysines identified did not (**Fig. 1D**). Overall, our results suggest that the lysine K686 of BCL11b is important for the global acetylation of this transcription factor.

### The lysine K686 of BCL11b regulates transcriptional activity of the p21 and IL-2 promoters

Our laboratory has previously demonstrated that the cellular factor BCL11b is recruited through Sp1 binding sites to the cyclin-dependent kinase inhibitor (CDKN1A/p21) promoter and that BCL11b silences *p21* gene transcription^3^. In CD4+ T lymphocytes and following interaction with p300, BCL11b has been shown to activate the IL-2 promoter following T-cell receptor (TCR) activation^7^. By mass spectrometry analysis, we identified that BCL11b was acetylated on its K686 lysine residue in unactivated condition (data not shown). To further study the impact of BCL11b acetylation on its transcriptional function, we performed luciferase reporter assays in the context of transient transfections of reporter constructs containing either the p21 or the IL-2 promoter cloned upstream of the luciferase reporter gene (**Fig. 2A and 2B**). These reporter constructs were co-transfected with increasing amounts of an expression vector for wild-type BCL11b or for a mutated BCL11b (K686R). We observed that expression of increasing amounts of BCL11b reduced p21 promoter activity, while increasing IL-2 promoter activity (**Fig. 2A, panels 2-4**) in agreement with previous studies^3,7,15^. Interestingly, when overexpressing the non-acetylated mutant K686R, we observed that both the repressive and activating transcriptional functions of BCL11b on the p21 and IL-2 promoters, respectively, were abolished (**Fig. 2A and 2B, panels 5-7, respectively**). These findings suggest that acetylation of lysine K686 of BCL11b is critical for its transcriptional function and that modulation of BCL11b acetylation state could be a potential therapeutic strategy for the treatment of diseases associated with dysregulated *CDKN1A/p21* or *IL-2* gene expression.

**Figure 2:**
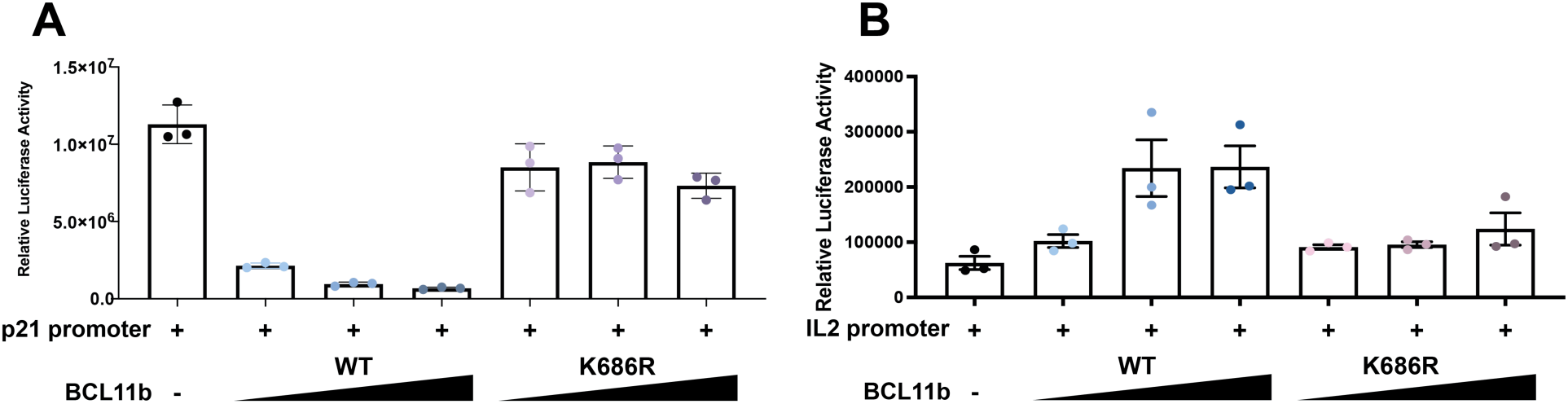
BCL11b K686R impacts the BCL11b-dependant transcriptional regulation of the p21 and IL-2 promoter. (A-B) HEK293T cells were transfected with the p21 **(A)** or the IL-2 **(B)** promoter -Luc reporter constructs in presence of increasing doses (0 ng, 200 ng, 400 ng, 800 ng) of BCL11b wild-type or mutated (K686R). To maintain the same amount of transfected DNA and to avoid squelching artifacts, the different amounts of BCL11b wild-type or mutated (K686R) cotransfected were complemented to 800 ng of DNA by using the corresponding empty plasmid. Forty-eight hours post-transfection, cells were collected, washed and lysed. Luciferase activity was measured from cellular lysates and normalized by protein dosage. Results are presented as histrograms. One representative experiment out of three is presented.

### The lysine K686 of BCL11b impacts its nuclear localization

Acetylation is known to be involved in the control of cytoplasmic and nuclear localization of non-histone proteins^21,22^. This prompted us to investigate whether lysine K686 might be involved in the nuclear localization of BCL11b. To do so, we performed a bioinformatic analysis using the NLS-mapper software, NLS-mapper (http://nls-mapper.iab.keio.ac.jp/ cgi-bin/NLS_Mapper_form.cgi; Kosugi et al., 2008, 2009), which identifies putative nuclear localization sequences (NLSs) within proteins (**Fig. 3A**). This analysis allowed us to identify a putative NLS sequence (in red) that includes lysine K686 (in blue) in BCL11b, with a cut-off score set to 3 (**Fig. 3A**). The cut-off score indicates the relative NLS activity, with higher scores indicating exclusively nuclear distribution and lower scores indicating cytoplasmic distribution^24^.

**Figure 3:**
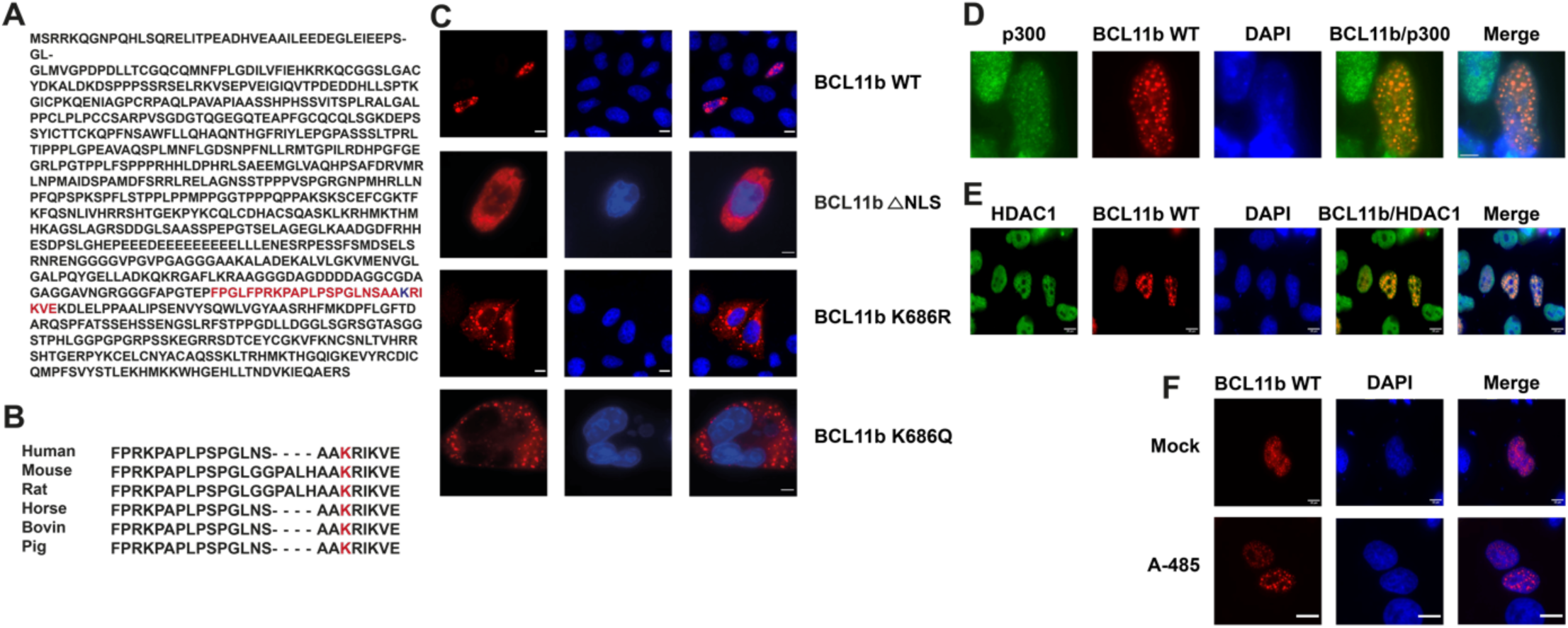
Lysine K686 of BCL11b is located in the NLS sequence of BCL11b and impacts the nuclear localization of BCL11b. **(A)** Schematic representation of our NLS mapper bioinformatic analysis which identified a putative NLS sequence in BCL11b (in red) containing the lysine K686 residue (in blue). **(B)** Sequence aligment between different species showing the putative NLS sequence identified in BCL11b. **(C)** HeLa cells were transfected with an expression vector either for wild-type BCL11b, either for BCL11b depleted in its putative NLS sequence, or for mutated forms of BCL11b (K686R, K686Q). Forty-eight hours post-transfection, cells were fixed, washed and subjected to observation under a fluorescence microscope. Scale bars are set to 10 μm. **(D-E)** HeLa cells were transfected with the expression vector for the wild-type BCL11b protein and with an expression vector for either HDAC1 or p300. Forty-eight hours post-transfection, cells were fixed, washed and subjected to observation under a fluorescence microscope. Scale bars are set to 10 μm. **(F)** HeLa cells were transfected with an expression vector for the wild-type BCL11b protein. Cells were treated for 16 h with 10 µM of A-485. Forty-eight hours post-transfection, cells were fixed, washed and subjected to observation under a fluorescence microscope. Scale bars are set to 10 μm.

We also examined the conservation of the putative BCL11b NLS sequence and of lysine K686 in different species and found that they were highly conserved, suggesting a functionnal importance for BCL11b (**Fig. 3B**). To further examine the influence of the putative nuclear localization sequence (NLS) and of lysine K686 of BCL11b on its nuclear localization, we conducted immunofluorescence analysis using HeLa cells transiently cotransfected with different constructs: mCherry-fused wild-type BCL11b (mCherry-BCL11b), mCherry-fused BCL11b with the putative NLS sequence deleted (ΔNLS BCL11b), mCherry-fused BCL11b with a mutation of lysine K686 to arginine (BCL11b K686R) and mCherry-fused BCL11b with a mutation of lysine K686 to glutamine (BCL11b K686Q) which mimics an acetylated amino acid. Our results indicated that wild-type BCL11b was localized in the nucleus (**Fig. 3C, panel 1**), in agreement with previous reports^14,25^. However, deletion of the putative NLS sequence of BCL11b (ΔNLS BCL11b) or mutation of lysine K686 (BCL11b K686R and BCL11b K686Q) abolished its translocation from the cytoplasm to the nucleus (**Fig. 3C, panels 2-4**). These findings indicate that the putative BCL11b NLS sequence including lysine K686 plays a critical role in the nuclear localization of BCL11b. However, the acetylation of lysine K686 does not have any effect on the cellular localization of BCL11b.

We thus demonstrated that mutation of lysine K686 in BCL11b globally affected its acetylation, as shown by our mass spectrometry analysis indicating that no acetylated residue was observed once K686 was mutated. We hypothesized that global acetylation might influence the subcellular localization of BCL11b. To test this hypothesis, we transiently cotransfected a BCL11b expression vector into HeLa cells and modulated BCL11b acetylation status by overexpressing p300 or HDAC1 to force acetylation or deacetylation, respectively. We analyzed the localization of acetylated or deacetylated BCL11b by immunofluorescence. We observed that p300 was predominantly localized in the nucleus (**Fig. 3D, panel 1**), in agreement with previous reports^26,27^. In presence of p300, we observed that mCherry_BCL11b_WT was localized specifically to the nucleus (**Fig. 3D, panel 2**). Previous studies have demonstrated an interaction between p300 and BCL11b^6,28^, and we here confirmed their colocalization when both proteins were expressed (**Fig. 3D, panel 4**). Moreover, HDAC1 was localized in the nucleoplasm (**Fig. 3E, panel 1**) and deacetylation did not affect the nuclear localization of wild-type BCL11b, as shown in **Figure 3E (panel 2)**. We also observed colocalization of BCL11b and HDAC1 in the nucleus (**Fig. 3E, panel 4**), in agreement with previous reports^6,10,25^. Overall, our results indicate that overexpression of p300 or of HDAC1 did not affect the nuclear localization of wild-type BCL11b.

Next, we further studied the impact of BCL11b acetylation on its nuclear localization by using a specific acetyltransferase inhibitor A-485. A-485 is a small molecule that binds to the catalytic site of p300 and competes with acetyl-CoA^29,30^. We performed transient cotransfection of a BCL11b expression vector into HeLa cells and we treated the cells with the specific p300 inhibitor A-485, four hours post-transfection. Sixteen hours post-transfection, cells were collected and analyzed under a fluorescence microscope. BCL11b was localized in the nucleus (**Fig. 3E, Mock**). Our results showed no variation in the localization of wild-type BCL11b in the presence of the A-485 inhibitor (**Fig. 3E, A-485**).

Taken together, our results demonstrate that the acetylation or deacetylation of BCL11b in the presence of p300 or of HDAC1, respectively, do not impact the nuclear localization of BCL11b.

### Mutation of lysine K686 in BCL11b impacts its interaction with p300

BCL11b is known to interact with various cellular factors, allowing it to positively or negatively regulate transcription of cellular or viral genes^31^. We therefore hypothesized that mutation of lysine K686 in BCL11b could impact its interaction with cellular partners. To test this hypothesis, we performed transient cotransfections of HEK293T cells with expression vectors for wild-type BCL11b or for K686R mutated BCL11b. At fourty-eight hours post-transfection, we immunoprecipitated BCL11b followed by western blot analysis using specific antibodies to determine whether the K686 lysine residue in BCL11b could impact its protein-protein interactions.

We successfully immunoprecipitated the wild-type and K686R mutated BCL11b proteins, and subsequent western blot analysis against various interacting partners revealed that the K686R mutation specifically affected its interaction with the cofactor p300 (**Fig. 4A**), while other cellular factors were not affected by this K686R mutation (**Supplementary data Fig. S1**). To further investigate the BCL11b-p300 interaction, we performed immunofluorescence experiments to determine whether the loss of interaction between p300 and the K686R mutated BCL11b was due to the cytoplasmic localization of BCL11b. We transiently cotransfected HeLa cells with a p300-Myc tagged expression vector and with the mCherry_BCL11b_K686R expression vector. We observed that, despite the cytoplasmic localization of K686R BCL11b and of p300, both proteins did not colocalize (**Fig. 4B**), in agreement with our immunoprecipitation experiments. These findings indicate that the mutation in lysine K686 of BCL11b impacts the interaction of BCL11b with p300 independently of the cytoplasmic localization of the K686R mutated BCL11b.

**Figure 4:**
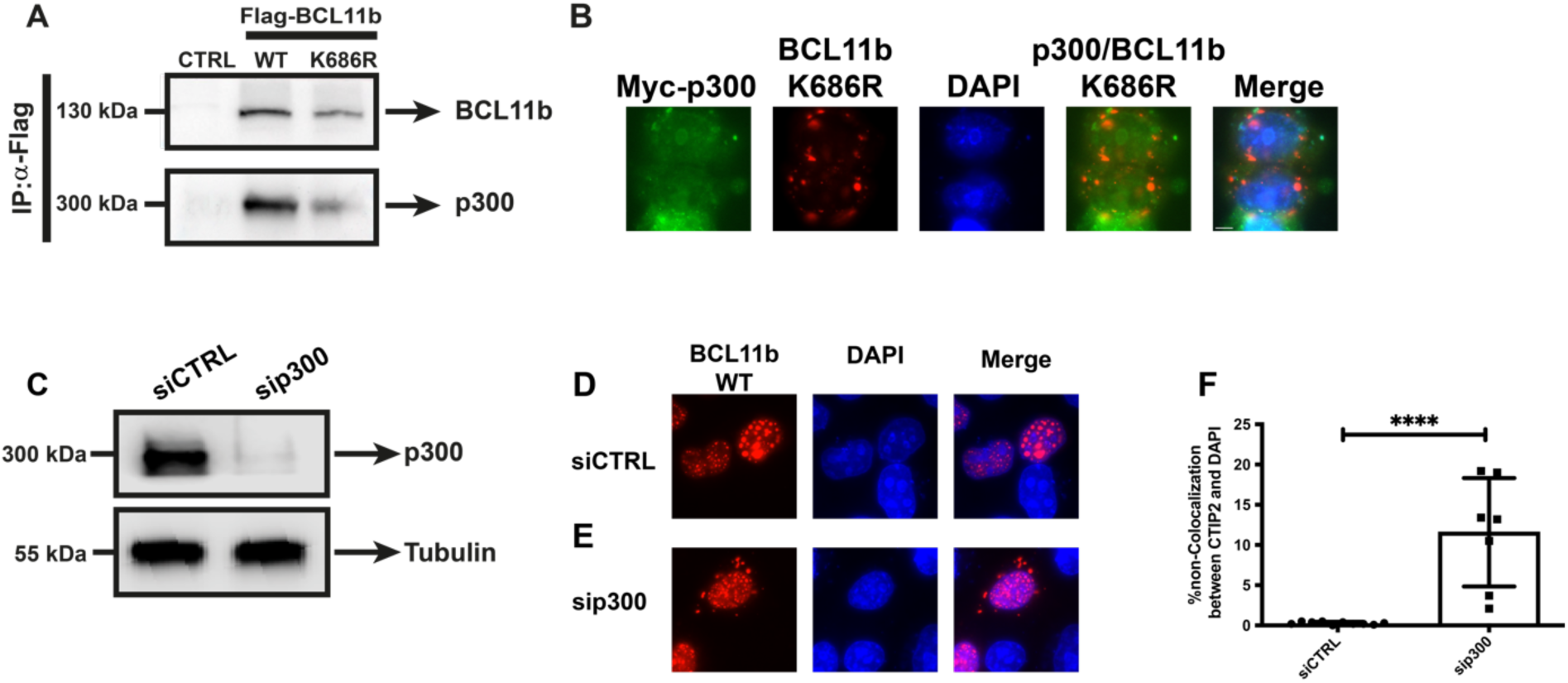
The acetyltransferase p300 impacts the nuclear localization of BCL11b. **(A)** HEK293T cells were transfected with an expression vector for either wild-type BCL11b or mutated (K686R) BCL11b, or with the corresponding parental empty vector (CTRL) as negative. Forty-eight hours post-transfection, the total protein extracts were collected and immunoprecipitated with an antibody against Flag-BCL11b or with IgG as negative control. The eluted protein complexes were analyzed by Western blot using specific antibodies against BCL11b or p300 **(B).** HeLa cells were transfected with the expression vector for mutated (K686R) BCL11b and with the expression vector for the tagged Myc-p300 protein. Forty-eight hours post-transfection, cells were fixed, washed and subjected to observation under a fluorescence microscope. Scale bars are set to 10 μm. **(C-E)** HeLa cells were transfected with the non-targeting siRNA (siCTRL) or with the p300 siRNA (si-p300). Forty-eight hours post-transfection, HeLa cells were transfected with an expression vector for the wild-type BCL11b. **(C)** Total protein extracts were collected 24 hours after the last transfection and were analyzed by Western blot using specific antibodies against p300 or tubulin as control. **(D-F)** Twenty-four hours after the last transfection, cells were fixed, washed and subjected to observation under a fluorescence microscope. Nuclei were counterstained by DAPI. Scale bars are set to 10 μm. **(F)** Inverse Manders correlation was performed to quantify the cytoplasmic BCL11b. A T-test was performed on 7 cells from 3 independent experiments (P****<0,0001).

### Downregulation of p300 impacts the nuclear localization of BCL11b

We just showed that mutation in lysine K686 of BCL11b impacted its interaction with p300 and its nuclear localization. Furthermore, although p300 is known to function as an acetyltransferase, studies have demonstrated a role of p300 as a nuclear transporter for the transcription factors SMAD2/3 (nuclear translocation of mothers against decapentaplegic homolog 2/3) and TAZ (also called WWTR1, WW domain-containing transcription regulator protein 1) in hepatic stellate cells stimulated with transforming growth factor B^32^. Based on this, we hypothesized that p300 may act as a nuclear transporter of BCL11b. To assess this, we employed RNA interference to induce depletion of endogenous p300 in HeLa cells. HeLa cells were transfected with several siRNAs targeting p300 mRNA (si-p300) or with a control non-targeting siRNA (siNT). Knockdown of p300 was validated by western blot (**Fig. 4C**) and immunofluorescence (**Supplementary Fig. S2**). Next, we evaluated by fluorescence microscopy HeLa cells expressing the mCherry_BCL11b_WT. We observed that BCL11b localized in the nucleus using HeLa cells transfected with the mCherry_BCL11b_WT and with the non-targeting control siRNA (**Fig. 4D**). Interestingly, depletion of p300 by RNA interference induced a partial translocation of mCherry-BCL11b to the cytoplasm (**Fig. 4E**). In order to quantify the intracellular abundance of BCL11b within the cytoplasm, we employed the inverse Manders coefficient to quantify the co-localization between the mCherry-BCL11b protein and the nuclear marker DAPI. Our results reveal a significant disparity in the cytoplasmic distribution of BCL11b between the siRNA targeting p300 and the non-targeting control siRNA. More specifically, our findings demonstrate a marked reduction in the cytoplasmic presence of BCL11b upon siRNA-mediated downregulation of p300, in comparison to the non-targeting control siRNA, suggesting that p300 might be involved in the cellular localization of BCL11b (**Fig. 4F**).

## Discussion

Acetylation is a post-translational modification catalyzed by acetyltransferases that add an acetyl group to internal lysine residues in eucaryotic proteins. Acetylation of lysine residues in histones neutralizes their positives charges and alters the size of their lysine lateral chains, inducing a modification of the protein conformation^21,33,34^. While the SUMOylation and phosphorylation of BCL11b have been previously studied in the literature^7,15,18,19^, acetylation of BCL11b had not been studied so far. In the present study, we demonstrated that BCL11b was acetylated by the acetyltransferase p300 at different lysine residues (K434, K585, K597, K686, K802, K851). We also found that mutation of lysine K686 to arginine, which mimics a non-acetylated and unacetylatable lysine, affected the global acetylation of BCL11b. Dubuissez and colleagues have previously shown that phosphorylation of serine 2 in the N-terminal domain of BCL11b abolishes its interaction with MTA1, which is associated with BCL11b to repress the IL-2 and Id-2 promoters^15^. Additionally, SUMOylation at lysine 679 in the C-terminal domain of BCL11b enables the recruitment of p300 to activate transcription^15^. These findings suggest that post-translational modifications of BCL11b can regulate both its repressive and activating functions in transcription. BCL11b has been shown to act as a transcriptional repressor of the p21 promoter and as an activator of the IL-2 promoter^3,15^. In addition to the impact on BCL11b global acetylation, we showed here that the BCL11b K686R mutation also had an impact on the functional role of BCL11b. Indeed, we showed that the lysine K686 mutation affected both the repressive role of BCL11b on the p21 promoter and its activating role on the IL-2 promoter.

In addition, a bioinformatics analysis allowed us to determine that lysine K686 of BCL11b was located within the NLS sequence and that this residue is highly conserved across different species, indicating that it is crucial for BCL11b protein function. Our analyses are in agreement with recent findings from the group of Christian Schmidt showing that the BCL11b KRIKV motif meets the consensus criteria for monopartite-type 2 NLS (K(K/R)X(K/R))^35^. Furthermore, this latter study has reported that the KRIKV motif containing lysine K686 is essential for the nuclear localization of the BCL11b protein^35^. As acetylation has also been shown to affect protein localization^36^, we generated BCL11b recombinant mutants mimicking an acetylated or a deacetylated lysine by substituting lysine K686 with a glutamine or a arginine residue, respectively, while maintaining the amino acid positive charge. Our results showed that the mutation of lysine K686 in BCL11b induced cytoplasmic localization of the protein, which was in agreement with the findings from the group of Christian Schmidt^35^. Our results also demonstrated that the substitution of lysine K686 with a glutamine residue resulted in cytoplasmic localization of the BCL11b protein. Therefore, we demonstrated that only acetylation of the K686 residue did not impact the nuclear localization of BCL11b. Moreover, we showed that the BCL11b K686R mutation affected the global acetylation of the BCL11b protein. Altogether, we studied the impact of global acetylation on BCL11b localization and we showed that BCL11b acetylation did not affect its localization.

BCL11b is known to be a nuclear protein that interacts with numerous factors involved in transcriptional regulation of many cellular and viral genes. Given that we showed that mutation of lysine K686 in BCL11b was involved in its nuclear localization, we investigated the impact of this mutation on interaction with known BCL11b partners. Interestingly, we showed that p300 was no longer able to interact with the K686R BCL11b mutant. p300 is mainly located in the nucleus, but it has also been demonstrated to shuttle between the cytoplasm and the nucleus^32,37–39^. In the present study, we showed that depletion of p300 using RNA interference induced partial cytoplasmic localization of BCL11b, highlighting the importance of the interaction between BCL11b and p300. However, BCL11b was not completely relocalized to the cytoplasm. It is worth noting that Christian Schmidt’s team has shown that the KRIKV motif of BCL11b allows interaction with importin α KPNA2, which enables nuclear import of BCL11b^35^ and that we showed here that the mutation in lysine K686 of BCL11b did not affect the interaction with KPNA2 (**Supplementary Fig. S1**). Therefore, we hypothesized that BCL11b in the cytoplasm could interact with p300, allowing the shuttling of BCL11b between the cytoplasm and the nucleus. Additionally, KPNA2, through the BCL11b KRIKV motif, allows nuclear import of BCL11b.

Together, our results demonstrate for the first time that BCL11b can be acetylated, and that this acetylation is mediated at least in part by the acetyltransferase p300. Furthermore, we showed that the lysine K686 of BCL11b is important not only for the global acetylation of the protein, but also for its functional role and nuclear localization. Finally, we identified the lysine K686 of BCL11b as important for its interaction with p300, and showed that depletion of p300 induced partial relocalization of BCL11b to the cytoplasm.

## Materials and methods

### Cell lines and cell culture

The human embryonic kidney cell line HEK293T (RRID:CVCL0063) was obtained from the AIDS Research and Reference Reagent Program (NIAID, iNIH) and the human cervical cancer cell line HeLa was obtained from the American Type Culture Collection (ATCC, Manassas, VA) and were maintained in Dulbecco’s modified Eagle medium (DMEM) (Life Technologies) supplemented with 10% fetal bovine serum, 50 U/ml of penicillin and 50 μg/mL of streptomycin (Life Technologies). In addition, 1mM of pyruvate sodium was added to maintain the HEK293T cell line. Cells were cultivated at 37°C in a 5% CO_2_ atmosphere. All cells were negative for mycoplasma (Sigma-Aldrich, MP0040A kit).

### Antibodies and reagents

Antibodies and other chemical reagents used in this work are listed in Supplementary **Table S1**

### Plasmid constructs

The following plasmids: pcDNA3, flag-BCL11b, Tap-BCL11b, have been described previously^4,6,9,14,40^. The pGL3-IL-2-Luc plasmid was obtained from Professor Yves Collette and the pGL3-p21-Luc was obtained from Prof. Stéphane Emiliani (Institut Cochin, Paris, France). The pCI_p300_WT, pCI_p300_ ΔHAT, pCI_pCAF_WT and pCI_pCAF_ ΔHAT constructs were obtained from Professor Yoshihiro Nakatani (National Institutes of Health, Bethesda, Marylan, USA)^41^. The pCMV_CBP was obtained from Professor Vincent Bourse (University of Liège, GIGA, Belgium).

Mutagenesis experiments were performed to induce specific mutations on the different acetylated lysine residues using the pcDNA3_flag_BCL11b plasmid as template (by the QuickChange Site-directed mutagenesis method, Stratagene). Different mutations were generated with specific primers containing the specific mutations through DNA polymerase Pfu enzyme. Primers are listed in **Supplementary Table S2**. Briefly, the parental plasmid was digested with the restriction enzyme DpnI. Then the *de novo* plasmid containing mutation was transformed into XL1 Blue bacteria at 37°C. Mutated constructs were fully sequenced after identification. All these mutated constructs were digested by XbaI (R0145L, New England Biolabs) and AleI (R0685L, New England Biolabs) and were cloned in the pcDNA3_flag_BCL11b_WT vector digested with the same enzymes.

The pCMV_mCherry_BCL11b_WT, pCMV_mCherryBCL11b_K686R, pCMV_mCherry_BCLL11b_K686Q and pCMV_mCherry_BCL11b_ΔNLS vectors were constructed from pCMV_mCherry. Different fragments of BCL11b were generated by PCR with specific primers. Primers are listed in **Supplementary Table S2.** The parental plasmid was digested with the restriction enzymes EcoRI (R3101L, New England Biolabs) and Xho1 (R0146L, New England Biolabs). Specific amplified PCR fragments and digested parental plasmid were ligated with NEBuilder HiFi DNA Assembly (New England BioLabs; E2621).

### Transfection and mass spectrometry

HEK293T cells were seeded at 9 x 10^6^ cells per well, grown in a 15cm^2^ dish and were transfected using the calcium phosphate coprecipitation method with the pcDNA3_Flag_BCL11b_WT or pCDNA3_Flag_BCL11b_mutated constructs and with pcDNA_p300. Fourty-eight hours post-transfection, the cells were lysed on ice for 10 min using a buffer containing 50mM Tris-HCl pH 4.8, 150mM NaCl, 5mM EDTA, 0.5% NP-40. After 10 minutes, total lysate was centrifugated at 13,000 *g*, the supernatant was stored for Western blot analysis. Antiproteases were systematically added to the lysis buffers.

Immunoprecipitations were performed on 2000 μg of total protein extracts with 5 µg of the indicated antibody to target specific proteins overnight at 4°C under rotation. The day after, total protein extracts were incubated with 40 µl of Protein A Sepharose CL-4B beads (Santa Cruz Biochemical) for 1 hour at 4°C under rotation. After three washes with buffer containing 50 mM Tris-HCl pH 8, 150 mM NaCl, 5 mM EDTA and 0.5% NP-40, dried beads were resuspended in NH4HCO3 50mM buffer. Protein samples were then reduced with 10 mM DL-dithiothreitol for 45 minutes at 56°C and alkylated with 55 mM iodoacetamide for 30 minutes at room temperature in the dark. Proteins were then digested overnight with 1 µg of trypsin (Promega®, Madison, WI, USA) at 37°C.

Peptides were purified using Oasis HLB 3cc cartridge (Waters) according to the manufacturer protocol. Peptides were resuspended in 100% H_2_O/0.1% HCOOH and injected on a Triple TOF 5600 mass spectrometer (Sciex, Concord, Canada) interfaced to an EK425 HPLC System (Eksigent, Dublin, CA) and data were acquired using Data-Dependent-Acquisition (DDA). Peptides were injected on a separation column (Eksigent ChromXP C18, 150mm, 3µm, 120A) using a two steps acetonitrile gradient (5-25% ACN/0.1% HCOOH in 48 minutes then 25%-60% ACN/0.1% HCOOH in 20 minutes at 5 µl/min) and were sprayed online in the mass spectrometer. MS1 spectra were collected in the range 400-1250 m/z with an accumulation of 250 ms. The 20 most intense precursors with a charge state 2-4 were selected for fragmentation, and MS2 spectra were collected in the range of 50-2000 m/z with an accumulation of 100 ms; precursor ions were excluded for reselection for 12 s.

The generated .WIFF files were directly analysed by ProteinPilot software 4.5 (Sciex, Concord, Canada) and the recorded spectra were matched to peptides by the search algorithm Paragon (Sciex, Concord, Canada) using the database human Swiss-Prot part of UniProtKB [The UniProt]. Trypsin was selected as digestion enzyme. Oxidation at methionine was set as dynamic modification, carbamidomethylation as static modification at cysteine. Decoy database search was performed with target false discovery rate <1% (FDR).

### Total protein extracts and immunoprecipitation

HEK293T cells were seeded at a density of 4 × 10^6^ cells per well and grown in a 10 cm^2^ dish and the cells were transfected using the calcium phosphate coprecipitation method with the indicated vectors (18 μg). Forty-eight hours post-transfection, the cells were lysed on ice for 10 min using a buffer containing 50 mM Tris-HCl pH 4.8, 150 mM NaCl, 5 mM EDTA, 0.5% NP-40. After 10 minutes, total lysate was centrifugated at 13,000*g*, the supernatant was stored for Western blot analysis. Antiproteases were systematically added to the lysis buffers.

Immunoprecipitations were performed on 1000 μg of total protein extracts with the indicated antibodies to target specific proteins. Briefly, the total protein extracts were incubated with 5 µg of antibody overnight at 4°C under rotation. The day after, total protein extracts were incubated with 40µl of Protein A Sepharose CL-4B beads (Sigma; GE17-0780-01) for 1 hour at 4°C under rotation. After three washes with buffer containing 50 mM Tris-HCl pH 8, 150 mM NaCl, 5 mM EDTA and 0,5% NP-40, immunoprecipitated proteins were eluted by heating at 100°C during 5 minutes in 1 x Laemmli Sample Buffer. Immunoprecipitated protein complexes were subjected to Western blot analysis.

### Western blot

Western blotting was performed using 20 µg of total protein extracts. Total protein extracts were separated by 8% SDS-PAGE. Then, total protein extracts were transferred onto 45 µM nitrocellulose membrane. Immunodetection was performed using primary antibodies targeting the protein of interest. Horseradish peroxidase (HRP)-conjugated secondary antibodies were used for chemiluminescence detection (Cell Signaling Technology).

### Transient transfections and luciferase assays

HEK293T cells were seeded at 0.25×10^6^ cells per well and grown in a 12-well plate. The cells were cotransfected with 600 ng of reporter constructs and with increasing amount (200 ng to 800 ng) of either the pcDNA3_BCL11b_WT or the pcDNA3_BCL11b_K686R vector by the calcium phosphate transfection method according to the manufacturer’s protocol (Takara). Forty-eight hours post-transfection, cells were lysed, and luciferase activities were measured using the Single Glo luciferase reporter assay (Promega). Results were normalized for transfection efficiency using total protein concentrations.

### Fluorescence microscopy

HeLa cells were seeded at 0.8×10^6^ cells per well and grown on coverslip in a 12-well plate. Twenty-four hours later, the cells were transfected with the indicated vectors with the Turbofect transfection reagent (R0533; ThermoFisher), according to the instructions of the supplier. Forty-eight hours post-transfection, the cells were fixed with PBS containing 4% formaldehyde for 15 minutes. Then, cells were permeabilized for 10 minutes with PBS 1x containing 0.5% Triton. Primary antibodies were incubated for 1 hour. After three washes, secondary antibodies were incubated for 1 hour. The coverslip was mounted on a slide using the Mowiol (81381; Sigma) containing 1 μg/ml DAPI (D9542, Sigma). Cells fluorescence was observed under a fluorescence microscope and imaged using a Observer Z1 Zeiss microscope (Zeiss, Oberkochen, Germany) with a Zeiss Plan-Apochromat 100x/1.4 oil objective. Images were acquired using the microscope system software Axiovision and processed using Fliji.

### siRNA transfections/RNA interference experiments

HeLa cells were transfected with 10 nM of siRNA targeting p300 (DsiRNAs, Integrated DNA Technologies) with Lipofecatime RNAiMAX Reagent (12778100; ThermoFisher) or with 10n M of a control siRNA (siCTRL; 51011403; Integrated DNA Technologies) and cultured for 72 hours.

## Role of funding sources

Funding agencies providing financial support did not participate in the design, data analyses, interpretation, or writing of this study.

## Contributors

Conceived and designed the study: CVL. Conceived and designed the experiments: LV, SF and CVL. Performed the experiments: LV, CV, MG, SF, GR, EP, TL, MS, MBD, VM, MB, CS. Performed mass spectrometry experiments and anaysis: RW and DC. Analyzed and interpretated the data: LV, MG, CVL. Wrote the manuscript: LV and CVL. Supervised the study and acquired funding: CVL and OR. All authors read or provided comments on the manuscript. All data were validated by CVL and LV.

## Data sharing statement

All the data/analyses presented in the current publication will be made available upon request to the corresponding author.

## Declaration of interests

The authors declare no conflicts of interest.

## Acknowledgments

CVL acknowledges funding from the Belgian National Fund for Scientific Research (grant CR J.002117 F.R.S-FNRS, Belgium); the “Télévie” program of the F.R.S.-FNRS; the “Fondation Roi Baudouin”; the Internationale Brachet Stiftung (IBS); the Walloon Region (“Fonds de Maturation”); The “Amis des Instituts Pasteur à Bruxelles”, asbl.; the University of Brussels (ULB - Action de Recherche Concertée (ARC) grant); the French INSERM agency “ANRS/Maladies infectieuses émergentes”. The laboratory of CVL. is part of the ULB-Cancer Research Centre (U-CRC) (Faculty of Medicine, ULB). Work in OR’s laboratory was supported by grants from the French agency “ANRS/Maladies infectieuses émergentes” and the “Alsace contre le Cancer” Foundation. LV is a doctoral fellows from the Belgian “Fonds pour la formation à la Recherche dans l’Industrie et dans l’Agriculture (FRIA)” and next from the “Fondation Rose et Jean Hoguet”. MG is a postdoctoral fellow supported by “Les Amis des Instituts Pasteur à Bruxelles, asbl” and by the Marie Sklodowska Curie COFUND action. SF was a fellow of the PDR-program of the F.R.S.-FNRS and next of the Télévie-Program of the F.R.S.-FNRS. GR was a fellow of the Belgian ‘Fonds pour la Recherche dans l′Industrie et l′Agriculture’ (FRIA-FNRS) and of the Van Buuren Fundation (ULB). EP was a doctoral fellow from fellow from the “Télévie” program of the F.R.S.-FNRS and next from the “Fondation Rose et Jean Hoguet”. MS and MBD are doctoral fellows from the Belgian “Fonds pour la formation à la Recherche dans l’Industrie et dans l’Agriculture (FRIA)”. MB was a doctoral fellow from the Belgian “Fonds pour la formation à la Recherche dans l’Industrie et dans l’Agriculture (FRIA)” and next from “Les Amis des Instituts Pasteur à Bruxelles”, asbl. CVL is “Directrice de Recherches” of the F.R.S-FNRS.

---

**Supplementary figure S1:**
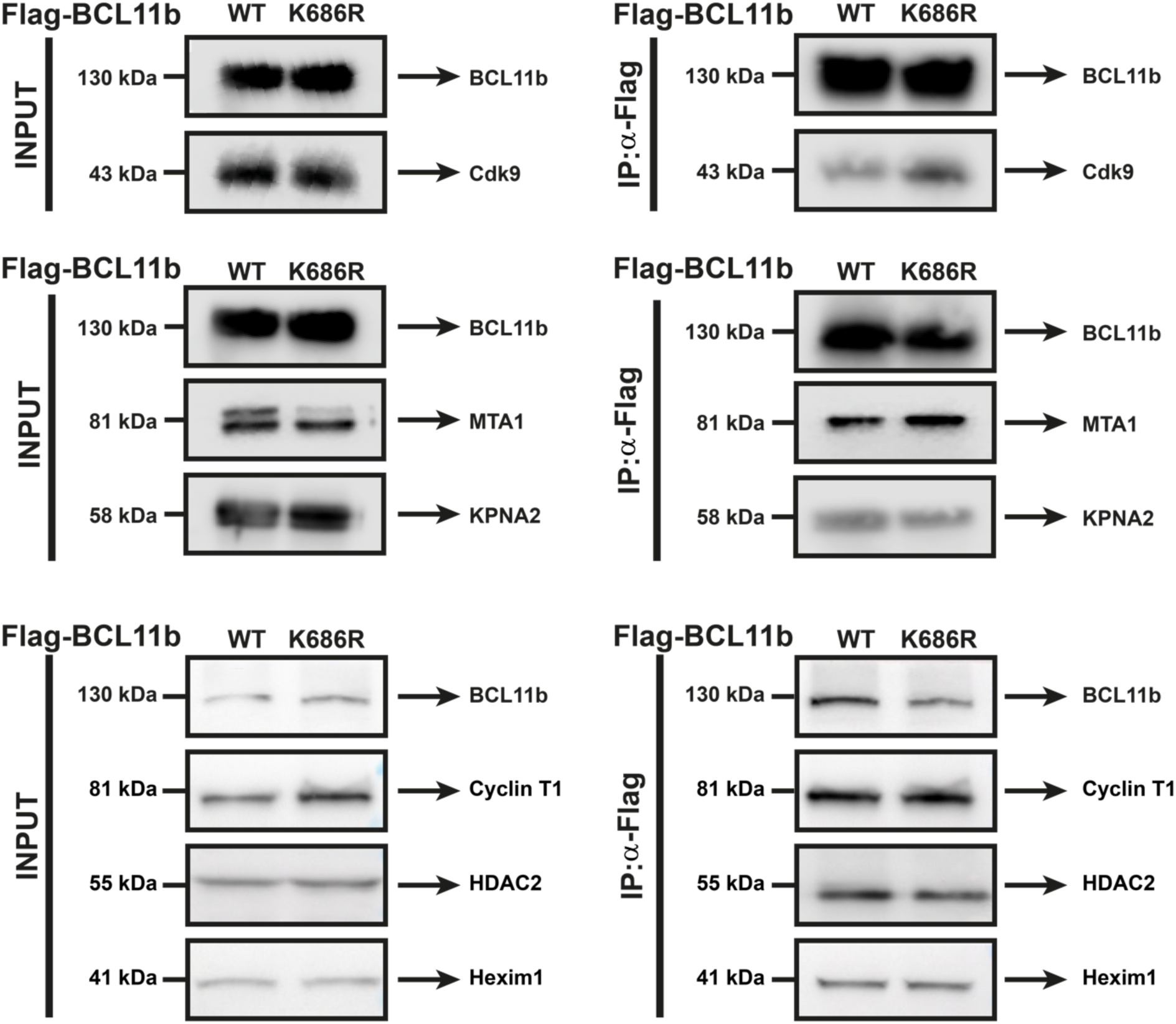
The K686R mutation in BCL11b does not impact the interaction with its partners. HEK293T cells were transfected with an expression vector for either the wild-type form of BCL11b or the mutated (K686R) form of BCL11b. Forty-eight hours post-transfection, total protein extracts were collected and immunoprecipitated with an antibody against Flag-BCL11b or with IgG as negative control. The eluted protein complexes were analyzed by Western blot using specific antibodies directed against BCL11b or against its associated cofactors as specified.

**Supplementary figure S2:**
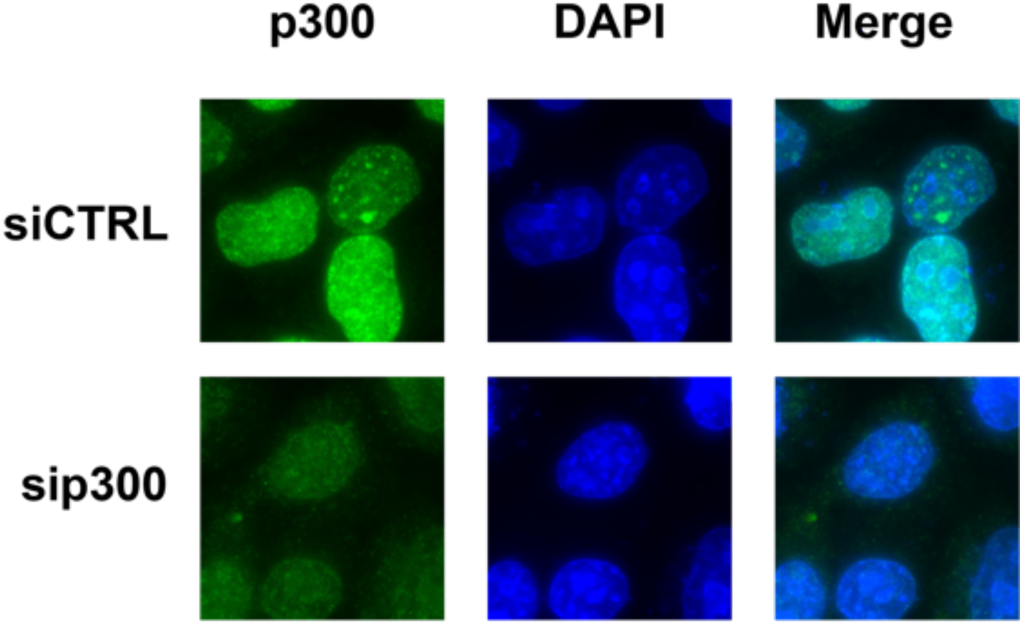
Knockdown of p300 impacts the nuclear localization of BCL11b. HeLa cells were transfected with a non-targeting siRNA (siCTRL) or with a targeting p300 siRNA (si-p300). Forty-eight hours post-transfection, HeLa cells were transfected with the expession vector for the wild-type BCL11b protein. Twenty-four hours after the last transfection, cells were fixed, washed and subjected to observation under a fluorescence microscope. Scale bars are set to 10 μm.

**Supplementary Table S1:** List of antibodies used.

| Antibodies | Reference |  |
| --- | --- | --- |
| IgG Rabbit | Cell Signaling | 2729S |
| IgG Mouse | Diagenode | C15400001 |
| Tubulin | Sigma | T5168 |
| Flag | Sigma | F3165 |
| KAc | UpState | 06-933 |
| p300 | Cell Signaling | 86377 |
| HDAC1 | Cell Signaling | 34589 |
| Myc | Cell Signaling | 2276S |
| HEXIM1 | Abcam | AB25388 |
| MTA1 | SantaCruz | SC-177773x |
| CDK9 | SantaCruz | Sc-13130X |
| KPNA2 | ProteinTech | 10819-1-AP |
| Cyclin T1 | SantaCruz | SC-8127 |
| HDAC2 | SantaCruz | SC-7899 |
| IgG Rabbit, HRP-linked | Cell Signaling | 7074 |
| IgG Mouse, HRP-linked | Cell Signaling | 7076 |
| Alexa Fluor 488 IgG Mouse | Thermofisher | A11001 |

**Supplementary Table S2:**
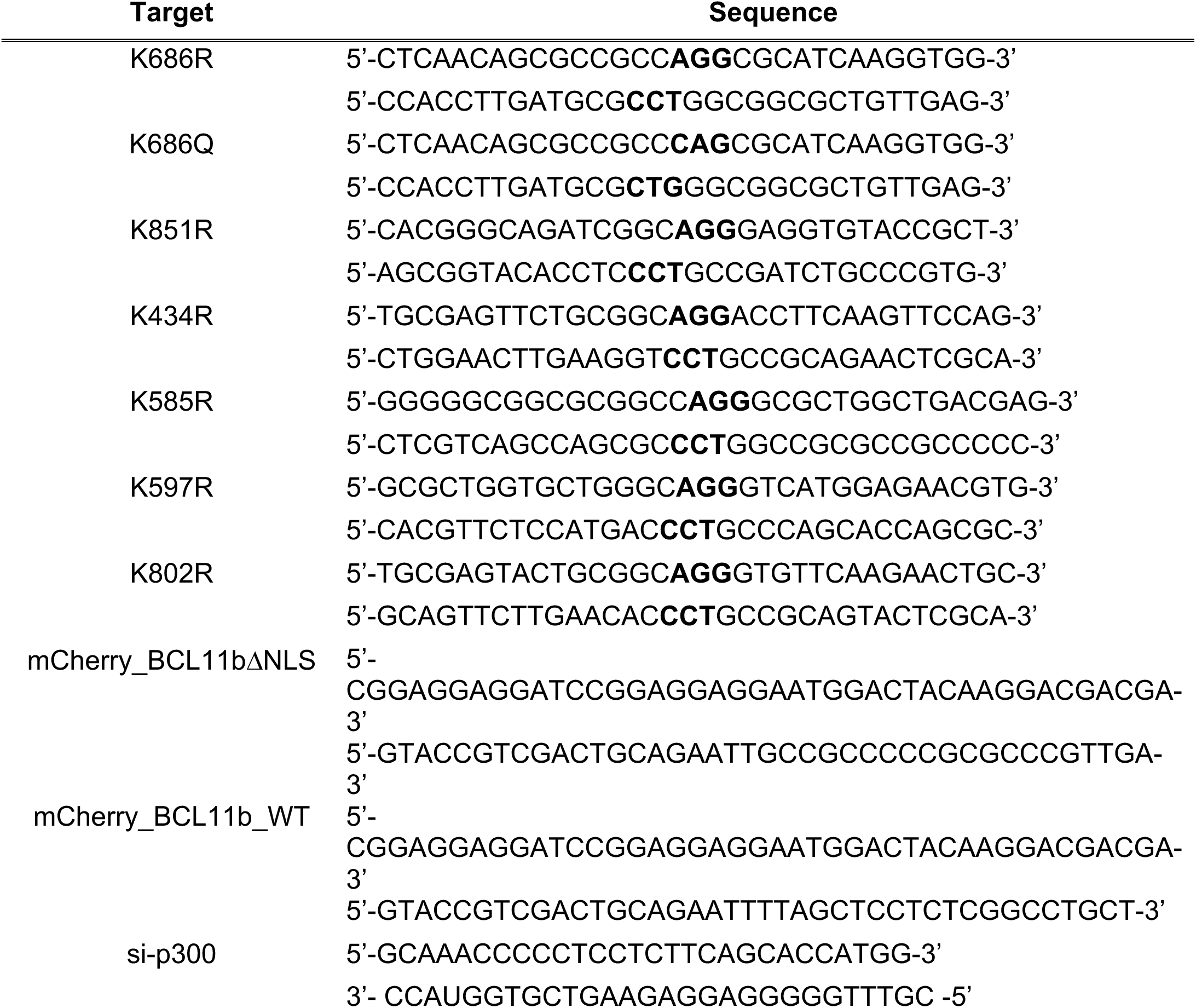
List of oligonucleotide sequences used. The introduced mutations are shown in bold.

## Notes

### Competing Interest Statement

The authors have declared no competing interest.

